# Recurrent SARS-CoV-2 spike mutations confer growth advantages to select JN.1 sublineages

**DOI:** 10.1101/2024.05.29.596362

**Authors:** Qian Wang, Ian A. Mellis, Anthony Bowen, Theresa Kowalski-Dobson, Riccardo Valdez, Phinikoula S. Katsamba, Lawrence Shapiro, Aubree Gordon, Yicheng Guo, David D. Ho, Lihong Liu

## Abstract

The recently dominant SARS-CoV-2 Omicron JN.1 has evolved into multiple sublineages, with recurrent spike mutations R346T, F456L, and T572I, some of which exhibit growth advantages, such as KP.2. We investigated these mutations in JN.1, examining their individual and combined effects on immune evasion, ACE2 receptor affinity, and in vitro infectivity. F456L increased resistance to neutralization by human sera, including those after JN.1 breakthrough infections, and by RBD class-1 monoclonal antibodies, significantly altering JN.1 antigenicity. R346T enhanced ACE2-binding affinity, without a discernible effect on serum neutralization. Individually, R346T and T572I modestly enhanced infectivity of each pseudovirus. Importantly, expanding sublineages such as KP.2 containing R346T, F456L, and V1104L, showed similar neutralization resistance as JN.1 with R346T and F456L, suggesting V1104L does not appreciably affect antibody evasion. Our findings illustrate how certain JN.1 mutations confer growth advantages in the population and could inform the design of the next COVID-19 vaccine booster.

## Introduction

SARS-CoV-2 continues to evolve and spread around the world, resulting in newly emerging variants with enhanced immune evasion, increased viral transmissibility, reduced vaccine efficacy, and other altered virological features. The Omicron JN.1 subvariant has been dominant, and prior studies have shown that it is already 2-3 times more resistant to serum neutralization than XBB.1.5^1,2^. JN.1 has also spawned multiple sublineages of related viruses, a number of which contains recurrent mutations in the spike protein, including R346T, F456L, and T572I (**Figure 1A and 1B**). These particular mutations are individually capable of enhancing antibody evasion (R346T and F456L), ACE2-binding and infectivity (R346T), or spike stability and conformation (T572I) when found on the genetic background of earlier Omicron subvariants^3–13^. Recent JN.1 subvariants with one or more of these recurrent mutations, such as KP.2 (R346T, F456L, and V1104L), appear to have a growth advantage, raising concern that they could become dominant by better evading antibodies raised by prior infections and vaccinations (**Figure 1C**).

**Figure 1.**
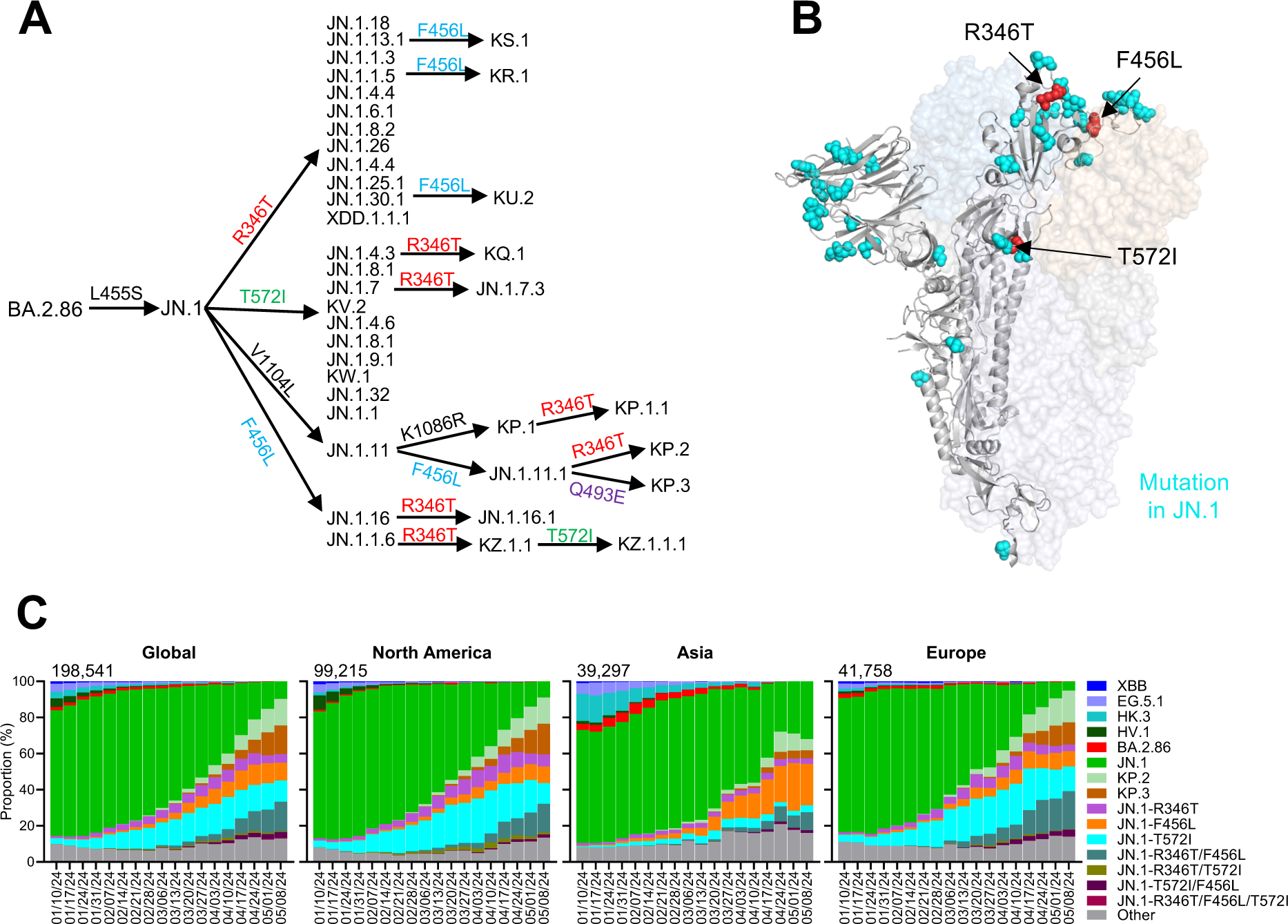
Recurrent spike mutations in SARS-CoV-2 Omicron JN.1 subvariants. **A.** Diversification in SARS-CoV-2 Omicron JN.1 subvariants with recurrent R346T, F456L, and T572I mutations up to May 8, 2024. **B.** Key spike mutations of JN.1 and its sublineage in the context of BA.2. JN.1 mutations relative to BA.2 are noted in cyan. Recurrent JN.1 sublineage mutations characterized in this study are noted in red. **C.** Frequency of SARS-CoV-2 variants, including JN.1 subvariants that carry the R346T, F456L, and T572I mutations. The value in the upper right corner of each box indicates the cumulative number of sequences for all circulating viruses within the specified time period.

A detailed understanding of the properties of recurrent JN.1 mutations could explain why certain sublineages are expanding and aid the design of future COVID-19 vaccine boosters. The latter scientific rationale is exemplified by the recent XBB.1.5 monovalent vaccine, where selecting a dominant and antigenically distinct spike protein for the vaccine formulation has led to improved serum neutralization of prevalent variants^1^. However, there have only been very limited studies reported on the antibody responses after JN.1 breakthrough infections^14–16^. Moreover, there is a lack of systematic evaluation of antigenicity of key JN.1 sublineages that carry various combinations of the aforementioned recurrent mutations.

Here, we present a comprehensive antigenic characterization of JN.1 sublineages featuring single, double, or triple recurrent mutations most frequently associated with growth advantages in the population. We investigated their ability to evade antibodies in sera from three clinical cohorts: those with XBB breakthrough infections or JN.1 breakthrough infections, as well as those with Omicron infections followed by a monovalent XBB.1.5 vaccine booster. Additionally, we assessed neutralization resistance of these subvariants to a panel of monoclonal antibodies targeting various epitopes of the viral spike protein. Furthermore, we measured their binding affinities to human ACE2 using two different assays and the infectivity of their pseudoviruses *in vitro*.

## Results

### Recurrent mutations in JN.1 sublineages associated with growth advantages

SARS-CoV-2 Omicron JN.1 had been dominant worldwide. Since January 2024, it has diversified into dozens of sublineages, many of which share recurrent spike mutations R346T (e.g., JN.1.18), F456L (e.g., JN.1.16), T572I (e.g., JN.1.7), or combinations of these mutations (e.g., KP.2) (**Figure 1A**). R346T and F456L are located in the spike receptor-binding domain (RBD), while T572I is in the SD1 domain of the spike protein (**Figure 1B**). Collectively, these recurrent mutations are found in approximately 77.3% of JN.1 samples globally. The prevalence of these JN.1 subvariants in Asia, Europe, and North America mirrors the global trend, with subvariants containing both F456L and R346T constituting over 16.1%, 38.6%, and 30.3% of cases in Asia, Europe, and North America, respectively (**Figure 1C**). The rapid increase of JN.1 sublineages that carry both the F456L and R346T mutations is exemplified by KP.2, which now accounts for over 14.7% of cases globally. Viruses with T572I increased during the early months of 2024. However, the frequency of T572I has been decreasing recently.

Notably, these recurrent mutations had previously been identified in several other SARS-CoV-2 variants. The R346T mutation was found in historically dominant strains such as BF.7, BA.4.6, BQ.1.1, XBB, EG.5, HK.3, and HV.1, whereas the F456L mutation was present in strains that dominated in 2023, such as XBB.1.16.6, EG.5, FL.1.5.1, HK.3, and HV.1. The T572I mutation was detected in several XBB subvariants, albeit at low frequencies^17^.

### Serum neutralization of JN.1 sublineages with recurrent mutations

To assess serum neutralization susceptibility of JN.1 sublineages, serum samples from 43 participants were collected from three clinical cohorts with a history of: 1) XBB breakthrough infection (XBB infx), 2) Omicron infection followed by XBB monovalent vaccine booster (Omicron infx + XBB.1.5 booster), or 3) JN.1 breakthrough infection (JN.1 infx). Demographic details along with vaccine and infection history for the three cohorts are summarized in **Table S1**. Individual details for each participant are included in **Table S2**. VSV-based pseudoviruses bearing subvariant spike proteins with all combinations of one, two, or three of the recurrent spike mutations R346T, F456L, and T572I, were generated and neutralization assays were conducted on these subvariants in parallel with ancestral D614G, as well as Omicron XBB.1.5 and JN.1.

Consistent with prior results, JN.1 was approximately 1.5-to-1.9 times as evasive to serum antibodies compared to XBB.1.5 in all three cohorts^1^ (**Figure 2A**). Among recurrent mutations, concordant results were observed across the cohorts. The F456L mutation alone led to serum neutralization titers 1.3-to 2.3-fold lower than JN.1. No substantial differences in titers were observed for mutation T572I or R346T. The different combinations of R346T, F456L, and T572I demonstrated various degrees of reduction in serum neutralizing titers compared with JN.1, but the reduction was statistically significant for any mutant virus that carried the F456L mutation (**Figure 2A**). Of note, JN.1 carrying all three recurrent mutations was most evasive to all sera tested (**Figures 2A and 2B**).

**Figure 2.**
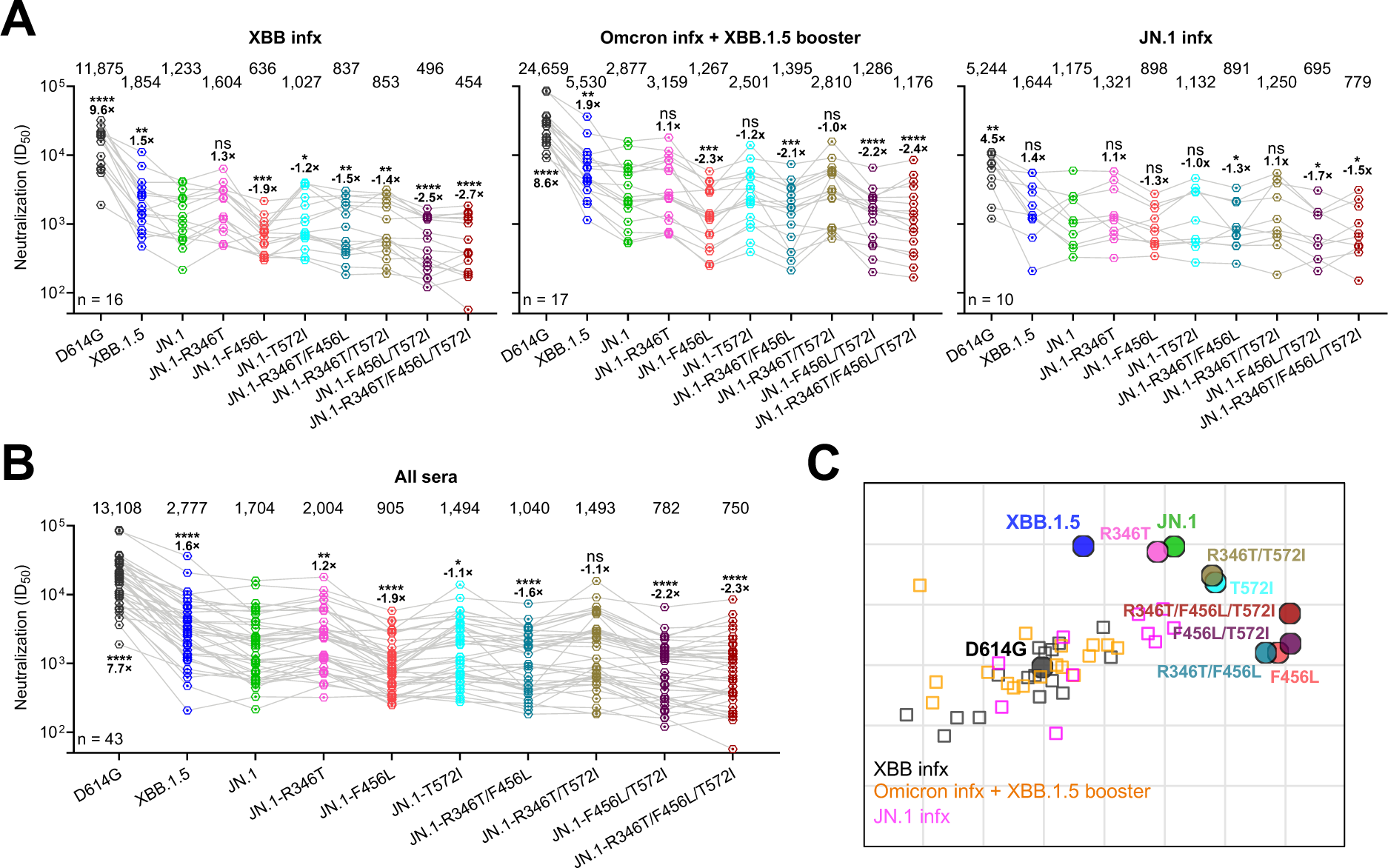
Serum neutralization of D614G, XBB.1.5, and JN.1 subvariants. **A.** Neutralizing ID_50_ titers of serum samples from “XBB infx”, “Omicron infx+XBB.1.5 booster” and “JN.1 infx” cohorts against the indicated SARS-CoV-2 variants. The geometric mean ID_50_ titers (GMT) are presented above symbols. Statistical analyses were performed by employing Wilcoxon matched-pairs signed-rank tests. n, sample size. ns, not significant; *p < 0.05; **p < 0.01; ***p < 0.001; ****p < 0.0001. **B.** Neutralizing ID_50_ titers of serum samples from all three cohorts in panel **A**. **C.** Antigenic map generated using neutralization data from panel **B**. D614G represents the central reference for all serum cohorts, with the antigenic distances calculated by the average divergence from each variant. One antigenic unit (AU) represents an approximately 2-fold change in ID_50_ titer. Serum samples and viruses are shown as squares and dots, respectively.

To examine overall patterns of antigenicity among the subvariants, we generated an antigenic map^18^ using all serum neutralization results (**Figure 2C**). The antigenic map reaffirmed the previously reported antigenic difference between JN.1 and XBB.1.5^1^. Sublineages of JN.1 harboring either the R346T mutation, the T572I mutation, or both, cluster closely with the parental JN.1 (within 1 antigenic unit), indicating similar antigenicity to the sera tested. In contrast, sublineages featuring the F456L mutation, regardless of the presence of R346T or T572I, form a distinct cluster with >1 unit more antigenic distance to D614G compared to JN.1. Additionally, F456L-containing sublineages were antigenically farther from XBB.1.5 than JN.1 (**Figure 2C**). These results indicate that F456L is the critical mutation driving the antigenic shift of select JN.1 sublineages. One such rapidly expanding progeny of JN.1 is KP.2, which contains R346T and F456L as well as V1104L in the S2 subunits of the spike. We specifically investigated its neutralization resistance to sera from those who had JN.1 breakthrough infections. Our data showed that KP.2 neutralization was similar or nearly identical to that of JN.1 with R346T and F456L (**Figure S1),** suggesting that V1104L has minimal impact on antibody neutralization.

### Monoclonal antibody neutralization of JN.1 sublineages

To better understand what types of neutralizing antibodies are impaired by these recurrent mutations on JN.1, the neutralization activity of a panel of 13 broadly neutralizing monoclonal antibodies (bnAbs) targeting known epitopes were evaluated. The panel included C1520^19^ and C1717^19^, which are directed to the N-terminal domain (NTD) and NTD-S2 interface, respectively, along with RBD-targeting antibodies such as S2K146^20^, Omi-18^21^, BD56-1854^22^, Omi-42^21^, S309 (sotrovimab)^23^, BD55-4637^22^, SA55^24^, 25F9^25^, 10-40^26^, and C68.61^27^. SD1-targeting antibodies were not included and only one RBD class 3 antibody (S309) was tested, as they were greatly impaired or completely knocked out by BA.2.86^28^, the immediate precursor of JN.1.

The half-maximal inhibitory concentration (IC_50_) values for each antibody against each pseudovirus are detailed in **Figure 3**. All these bnAbs retained activity against the D614G and XBB.1.5 variants. JN.1 demonstrated increased evasion against RBD class-1 antibodies, such as S2K146 and Omi-18, and the class-3 antibody S309, corroborating findings from previous studies^28,29^. JN.1 subvariants carrying R346T, T572I, or both, exhibited neutralization profiles similar to those of parental JN.1, aligning with our serum neutralization results (**Figure 2**). In contrast, any subvariant harboring the F456L mutation, alone or in combination, significantly impaired the neutralizing activity of RBD class-1 monoclonal antibodies BD-1854, BD57-1302, and Omi-42 (**Figure 3**).

**Figure 3.**
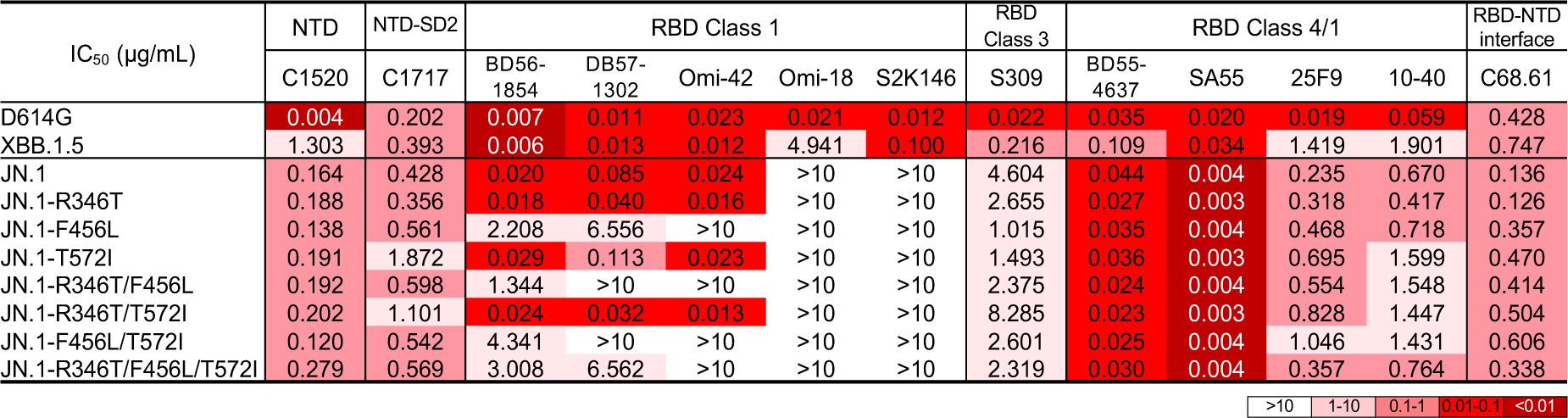
Resistance of D614G, XBB.1.5 and JN.1 subvariants to neutralization by monoclonal antibodies. The antibody concentrations resulting in 50% inhibition of infectivity (IC_50_) are presented.

The mutation T572I resides in SD1 domain of spike (**Figure 1B**), which could have an impact on antibodies directed to this domain. However, there are no published SD1-directed monoclonal antibodies that could neutralize JN.1^28,30,31^. We therefore assessed T572I in the genetic background of BA.2, which retains good sensitivity to neutralization by SD1-directed antibodies. Pseudovirus neutralization assays against BA.2 and BA.2-T572I were conducted using a panel of 15 bnAbs, including SD1-directed antibodies S3H3^32^ and C68.59^27^, and RBD/SD1-directed antibody 12-19^33^. BA.2-T572I was approximately 2-fold less sensitive to neutralization by the three additional monoclonal antibodies, SD1 or RBD/SD1, compared to BA.2 (**Figure S2A**). Conversely, BA.2-T572I was slightly more susceptible to neutralization by RBD class 1 and 4/1 antibodies, as well as to an antibody directed to the NTD/SD2. Admittedly, these differences are rather minor and unlikely to explain why sublineages containing T572I expanded transiently a few months ago. Structurally, T572I is not located within the epitope of mAbs S3H3 and 12-19, as shown in **Figure S2B**. This suggests that T572I influences the neutralization of these SD-1 antibodies through a conformational alteration of SD-1. Structural modeling indicates that while I572 can still be accommodated on the RBD of BA.2 in its downward conformation (**Figure S2C**), the T572I mutation disrupts the hydrogen bond between T572 and D568 on the BA.2 RBD in its upward conformation (**Figure S2D**). This disruption reveals that T572 may affect the dynamics of SD1 during the transition from the RBD down to the RBD up position.

### ACE2 affinity and infectivity of JN.1 sublineages with recurrent mutations

Another property that could confer a selective advantage to JN.1 sublineages is the affinity for the ACE2 receptor, which facilitates viral infectivity, and perhaps transmissibility as well. Therefore, we conducted surface plasmon resonance (SPR) measurement of the affinity of the full spike proteins for human ACE2 (hACE2) (**Figures 4A and 4B**), along with inhibition of JN.1 subvariants by soluble hACE2 (**Figure 4C**). A consistent finding was that the R346T mutation alone appeared to increase affinity for hACE2 by ∼1.5 fold. On the other hand, F456L and T572I did not discernibly change receptor affinity.

**Figure 4.**
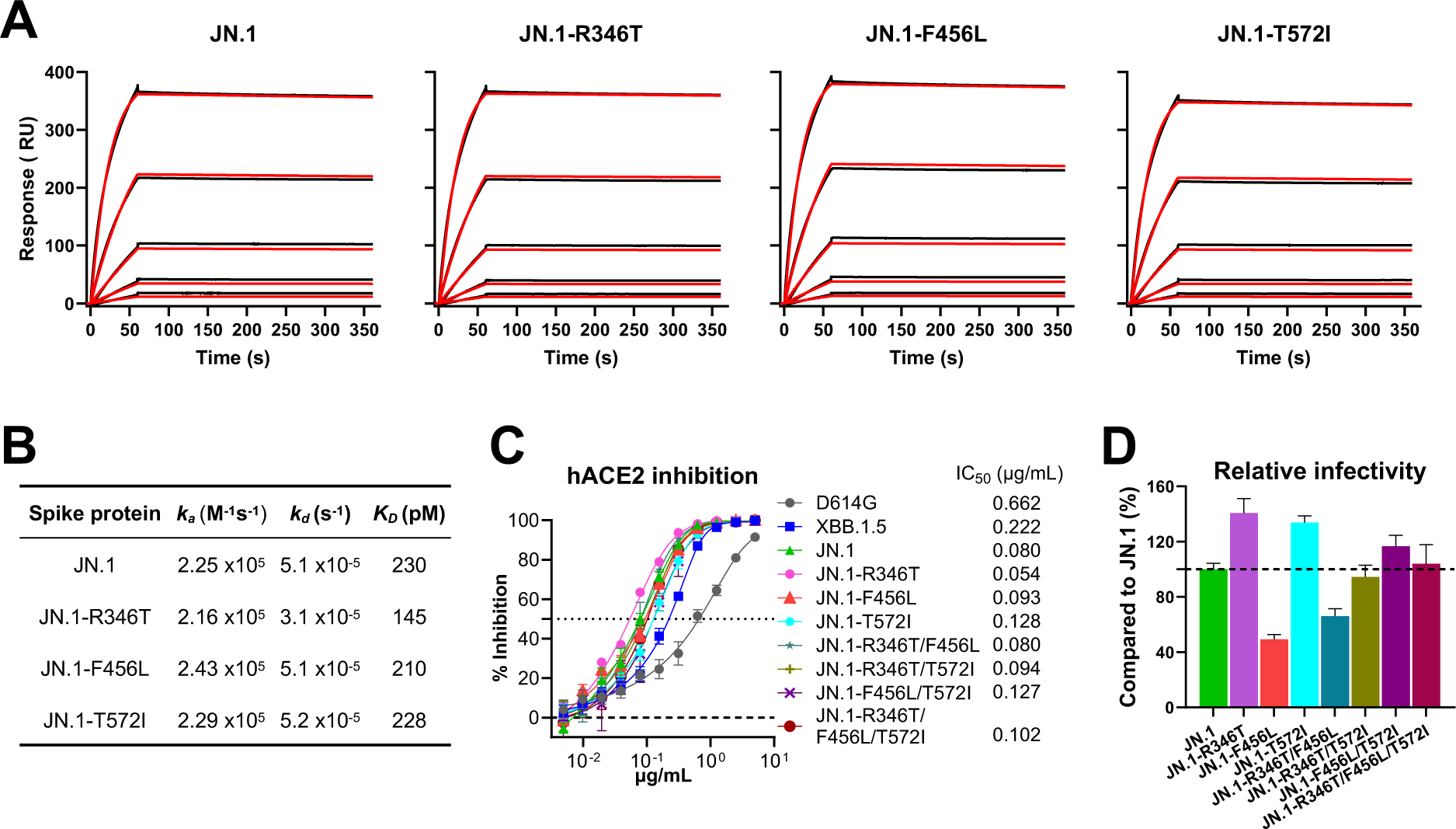
ACE2 affinity of JN.1 and its subvariants with individual spike mutation. A. Sensorgrams of the dose-dependent (200, 66.6, 22.2, 7.41, and 2.47 nM) binding curves (red lines) and the fitted data (black lines) of the indicated spike proteins to human ACE2 (hACE2) tested by surface plasmon resonance (SPR). B. Affinity (*K*_D_), association rate constant (*k*_a_) and dissociation rate constant (*k*_d_) of the indicated spike proteins to hACE2. C. Sensitivity of pseudotyped D614G, XBB.1.5, and JN.1 subvariants to hACE2 inhibition. IC_50_ values are also denoted. Data are shown as mean ± standard error of mean (s.e.m.) for four technical replicates. D. Relative infectivity of pseudotyped SARS-CoV-2 JN.1 subvariants compared to the parental virus JN.1 in Vero-E6 cells. Error bars represent standard error of the mean.

JN.1 infectivity of Vero-E6 cells in vitro was modestly increased by R346T or T572I, while it was somewhat reduced by F456L or the combination R346T+F456L. Other combinations of mutations did not markedly affect infectivity of Vero-E6 cells **(Figure 4D)**. The combination R346T+T572I did not noticeably alter infectivity, perhaps reflecting epistatic effects that can be further characterized in later studies.

## Discussion

In this study, we found that the predominant recurrent mutations R346T, F456L, and T572I confer a variety of phenotypic advantages to SARS-CoV-2 Omicron subvariant JN.1, providing an indication for why some of its progeny sublineages are becoming dominant in the population. R346T increases receptor binding affinity by ∼50% and pseudovirus infectivity by ∼40%, whereas T572I increases pseudovirus infectivity only slightly (**Figure 4**). However, most notably, F456L enhances antibody evasion (**Figure 2)** by impairing neutralization of antibodies directed to RBD class-1 region of the spike protein (**Figure 3**). R346T or T572I alone in the background of JN.1 has no detectable impact on serum neutralization (**Figure 2**). T572I in the background of BA.2 minimally impairs the neutralization of SD1-directed antibodies (**Figure S2A**), but their contribution to the total neutralizing activity of serum is likely very small. It is important to note the rapid expansion of the JN.1 sublineages, specifically KP.2 and KP.3. Each of these has a similar global prevalence, accounting for approximately 15% of the current JN.1 sublineages. KP.2 is characterized by the mutations R346T, F456L, and V1104L, whereas KP.3 exhibits F456L, Q493E, and V1104L. Remarkably, both dominant variants share the F456L mutation, again demonstrating that antibody evasion is a major selective advantage for viral growth in the population, as we and others have repeatedly observed over the past few years for successive variants and subvariants of SARS-CoV-2^1,8,34–36^.

Our findings serve as a reminder that SARS-CoV-2 has steadily become more and more antibody resistant, while overall serum antibody levels in the population continues to wane. Breakthrough infections will become more frequent with the passage of time, unless vaccine boosters are administered. The vaccine manufacturers, along with the governmental health agencies, are now contemplating the choice of the viral strain(s) to incorporate into the next updated COVID-19 vaccine^37,38^. Will the previously dominant JN.1 be selected, as the World Health Organization has recommended? Alternatively, will one of the rising JN.1 sublineages with F456L, such as KP.2 or KP.3 be chosen? A decision is expected within weeks. Our immunogenicity data presented herein could inform those deliberations, even though antigenicity does not directly reflect immunogenicity.

## Limitations

Limitations of this study include the small sample size, especially for the JN.1 breakthrough cohort. Testing sera from different clinical cohorts could enhance the general applicability of our findings. Additionally, the epistatic interactions among the recurrent spike mutations have not been addressed directly.

## ACKNOWLEDGEMENTS

This study was supported by funding from the NIH SARS-CoV-2 Assessment of Viral Evolution (SAVE) Program (subcontract no. 0258-A709-4609 under federal contract no. 75N93021C00014) and the Gates Foundation (project INV019355) to D.D.H., as well as by funding from NIH contract 75N93019C00051 and 75N93021C00016 to A.G., and internal startup funding from Columbia University to Y.G. We would also like to thank the Taikang Center for Life and Medical Sciences at Wuhan University, Wuhan, China, for providing funding support to L.L. We thank Zijin Chu, Carmen Gherasim, Anna Buswinka, Gabe Simjanovski, Joseph Wendzinski, Mayurika Patel, Kathleen Lindsey, Dawson Davis, for conducting the IASO and VIVA studies and Emily Stoneman, David Manthei, Victoria Blanc, Savanna Sneeringer, and Pamela Bennett-Baker for conducting the IASO study.

## AUTHOR CONTRIBUTIONS

L. L., A. G., and D. D. H. conceived and supervised the project. Q.W. managed the project. Q.W. and L.L. constructed the spike expression plasmids. Q.W., I.A.M., and L.L. conducted pseudovirus neutralization assays. Q.W. and L.L. purified SARS-CoV-2 soluble spike proteins, hACE2 protein and monoclonal antibodies. Y.G. conducted bioinformatic analyses. P.S.K. and L.S. generated the surface plasmon resonance (SPR) results. C.G., R.V., A.G., and A.B., provided clinical samples and organized information. Q.W., I.A.M., Y.G., A.B., L. L., and D.D.H. analyzed the results and wrote the manuscript. All authors have reviewed the results and have given their approval for the final version of the manuscript.

## Methods

### Resource availability

#### Lead contact

Further information and requests for resources should be directed to and will be fulfilled by the lead contact, Lihong Liu.

#### Materials availability

All requests for resources and reagents should be directed to and will be fulfilled by the Lead Contact Author. This includes selective cell lines, plasmids, antibodies, viruses, serum, and proteins. All reagents will be made available on request after completion of a Material Transfer Agreement.

#### Data and code availability Data

Data reported in this paper will be shared by the lead contact upon request.

#### Code

This paper does not report original code.

#### All other items

Any additional information required to reanalyze the data reported in this paper is available from the lead contact upon request.

### Experimental model and subjects

#### Sample collection

The sera samples were all collected at the University of Michigan through the Immunity-Associated with SARS-CoV-2 Study (IASO)^39^; XBB infx and Omicron infx + XBB.1.5 booster cohorts) and the VIVA Study (JN.1 cohort), and the collections were conducted under protocols reviewed and approved by the Institutional Review Board of the University of Michigan Medical School. All subjects provided written informed consent. Sera were collected from three cohorts:

Individuals who had an XBB sublineage infection (XBB infx) between February and September 2023; 2) Individuals with a prior Omicron sublineage infection between January 2022 and August 2023 followed by an XBB.1.5 monovalent vaccine booster (Omicron infx + XBB.1.5 booster); and 3) Individuals who had a JN.1 sublineage infection (JN.1 infx) in January or February 2024. Details for participants are described in **Tables S1 and S2**. All serum samples were heat inactivated at 56°C for 30 min before use.

#### Cell lines

Vero-E6 (CRL-1586) cells and HEK293T (CRL-3216) cells were obtained from ATCC and cultured at 37°C with 5% CO_2_ in Dulbecco modified Eagle medium (DMEM) + 10% fetal bovine serum (FBS) + 1% penicillin-streptomycin. Expi293 (A14527) cells were purchased from Thermo Fisher Scientific and maintained in Expi293 expression medium per the manufacturer’s instructions. Vero-E6 cells are derived from African green monkey kidneys. HEK293T cells and Expi293 cells are of human female origin.

#### Plasmid generation

As previously described, the antibody sequences for the heavy chain variable (VH) and the light chain variable (VL) domains were synthesized by GenScript, and then cloned into the gWiz vector to produce antibody expression plasmids. For the packaging plasmids for pseudoviruses, mutations were made by using the QuikChange II XL site-directed mutagenesis kit (Agilent) on the JN.1 construct that we previously generated^1^. For the soluble spike expression plasmids, in addition to the mutations under investigation, the 2P substitutions (K986P, V987P) and a “GSAS” substitution in the furin cleavage site (682-685aa) were introduced in the ectodomain (1-1208aa in WA1) of each of the spikes and then fused with an 8x His-tag at the C-terminus as previously described^40^. All constructs were verified using Sanger sequencing prior to use.

### Experimental Methods

#### Protein expression and purification

The gWiz-antibody, paH-spike, or pcDNA3-sACE2-WT(732)-IgG1 (Addgene plasmid #154104) plasmid^41^ was transfected into Expi293 cells using PEI at a ratio of 1:3, and then the supernatants were collected after five days. The antibodies and human ACE2 (hACE2) fused to a Fc tag were purified with Protein A Sepharose (Cytiva) following the manufacturer’s instructions. For SPR analysis, the hACE2 protein was further purified with Superdex 200 Increase 10/300 GL column. Spike proteins were purified using Ni-NTA resin (Invitrogen) per the manufacturer’s instructions. Molecular weight and purity were confirmed by SDS-PAGE protein electrophoresis prior to use.

#### Surface plasmon resonance (SPR)

SPR assays to assess hACE2 binding to SARS-CoV-2 spike proteins were conducted using a Biacore T200 biosensor, equipped with a Series S CM5 chip (Cytiva, Cat# BR100530), maintained at 25°C. The running buffer consisted of 10 mM HEPES pH 7.4, 150 mM NaCl, 0.2 mg/mL BSA, and 0.01% (v/v) Tween 20. The spike proteins were captured via their C-terminal His tag on an anti-His antibody surface, generated using the His-capture kit (Cytiva, Cat# 28,995,056), following the manufacturer’s instructions.

In each binding cycle, each spike was captured over individual flow cells at approximately 500-700 RU, with an anti-6×His antibody surface serving as the reference flow cell. hACE2 was prepared in five different concentrations using a three-fold dilution series in running buffer, ranging from 2.47 to 200 nM. Samples were sequentially tested with increasing protein concentrations. Blank buffer cycles were performed by injecting running buffer instead of the analyte, after two hACE2 injections. The association and dissociation rates were monitored for 90 seconds and 300 seconds respectively, at a flow rate of 50 µL/min. Bound spike/ACE2 complexes were removed using a 10-second pulse of 15 mM H3PO4 at 100 µL/min, enabling the regeneration of the anti-6×His surface for a new capture cycle of each spike, followed by a 60-second buffer wash at 100 µL/min. Data were analyzed and fitted to a 1:1 interaction model using Scrubber 2.0 (BioLogic Software).

#### Pseudovirus production

SARS-CoV-2 pseudoviruses were produced in a vesicular stomatitis virus (VSV) background, in which the native VSV glycoprotein was replaced by SARS-CoV-2 spike and its variants, as previously described^42^. Briefly, plasmids containing the appropriate spike were transfected into HEK293T cells with PEI. After 24 hours, VSV-G pseudotyped ΔG-luciferase (G*ΔG-luciferase, Kerafast) was added, and then washed with culture medium three times before being cultured in fresh medium for another 24 hours. Anti-VSVG (I1) antibody^43^ was added to deplete non-pseudotyped viruses. Pseudoviruses were then harvested, centrifuged, and then aliquoted and stored at −80°C.

#### Pseudovirus infectivity

Pseudovirus particles bearing various SARS-CoV-2 spike proteins were inoculated onto Vero-E6 cells, starting with 50 µl per well in 96-well plates and then subjected to serial dilutions. Following a 16–18-hour incubation at 37°C, we measured the activity of the virus-encoded firefly luciferase in the cell lysates. Subsequently, the luciferase activity for each tested pseudotyped virus, which was not oversaturated, was normalized to the parental D614G S control, with an infectivity set to 1.0.

#### Pseudovirus neutralization assays with sera, mAbs, or ACE2

Each SARS-CoV-2 pseudovirus was titrated to standardize viral infectious dose before use in neutralization assays. Serially diluted (seven dilutions of) heat-inactivated sera or antibodies were added in 96-well plates, starting at 1:100 dilution for sera and 10 µg/mL for antibodies. For ACE2 inhibition assays, as previously reported, we used soluble chimeric human ACE2, which contains ACE2 residues 1-732 fused to human IgG1 Fc. hACE2 was diluted starting from 10 µg/mL with a dilution factor of two across 11 serial dilutions. Then, pseudoviruses were added and incubated at 37 °C for 1 hour. In each plate, wells containing only pseudoviruses were included as controls. Vero-E6 cells were then added at a density of 4 × 10^4^ cells per well and incubate at 37 °C for an additional 16 hours. Cells were lysed and luminescence was determined by the Luciferase Assay System (Promega) and SoftMax Pro v.7.0.2 (Molecular Devices) according to the manufacturers’ instructions. Data were analyzed in GraphPad Prism v.9.3.

#### Antigenic cartography

Antigenic distances between sera, D614G, XBB.1.5, JN.1 and other SARS-CoV-2 JN.1 subvariants were determined by integrating all ID_50_ values of individual serum samples through a published antigenic cartography approach^18^. The visualization was generated using Racmacs (v.1.1.4, https://acorg.github.io/Racmacs/) in R version 4.0.3. The optimization step count was set at 2,000 and the minimum column basis parameter set to ‘none’, the ‘mapDistances’ function was employed to calculate antigenic distances between each serum sample and variant.

#### Quantification and statistical analysis

Neutralization ID_50_ and IC_50_ values were determined by fitting a five-parameter dose-response curve in GraphPad Prism v9.3. Statistical significance of differences in neutralizing titer was evaluated using two-tailed Wilcoxon matched-pairs signed-rank tests in GraphPad Prism v9.3. Significance is presented as following: ns, not significant; *p < 0.05; **p < 0.01; and ***p < 0.001, and ****p < 0.0001.

## Supplementary Information

**Figure S1.**
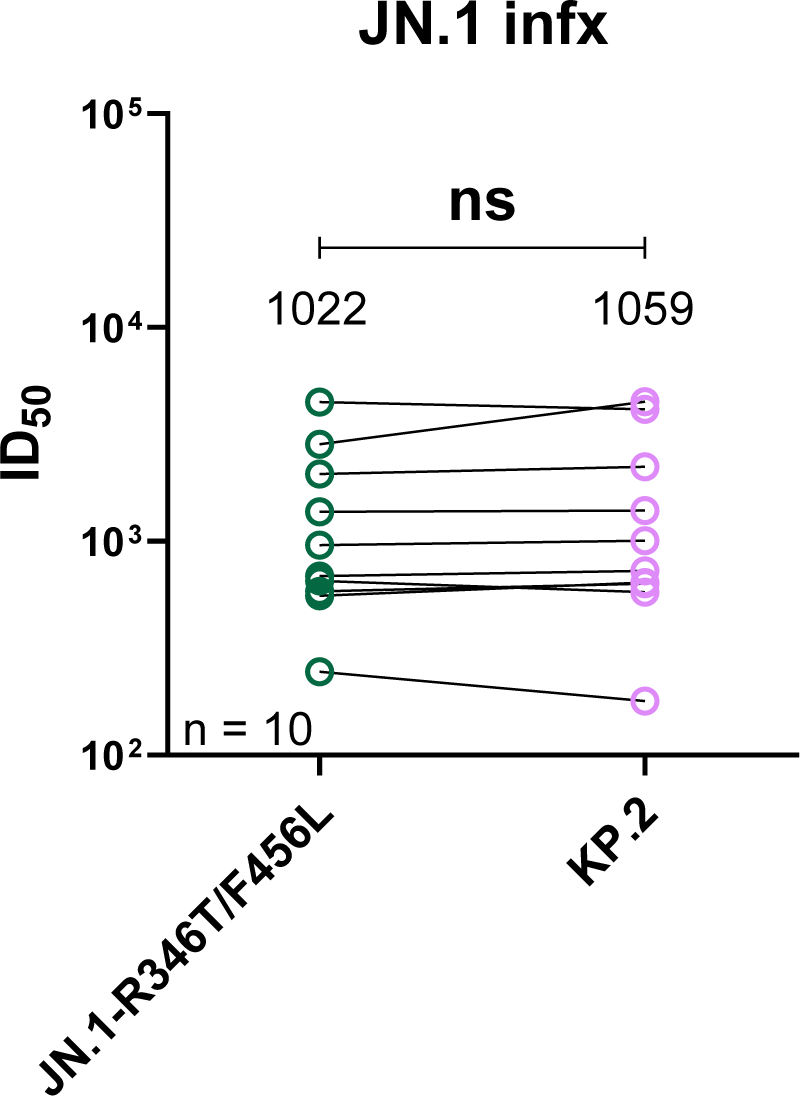
Serum neutralization of JN.1-R346T/F456L and KP.2. Neutralizing ID_50_ titers of serum samples from the “JN.1 infx” cohort against JN.1-R346T/F456L and KP.2. The geometric mean ID_50_ titers (GMT) are presented above symbols. Statistical analyses were performed by employing Wilcoxon matched-pairs signed-rank tests. n, sample size. ns, not significant.

**Figure S2.**
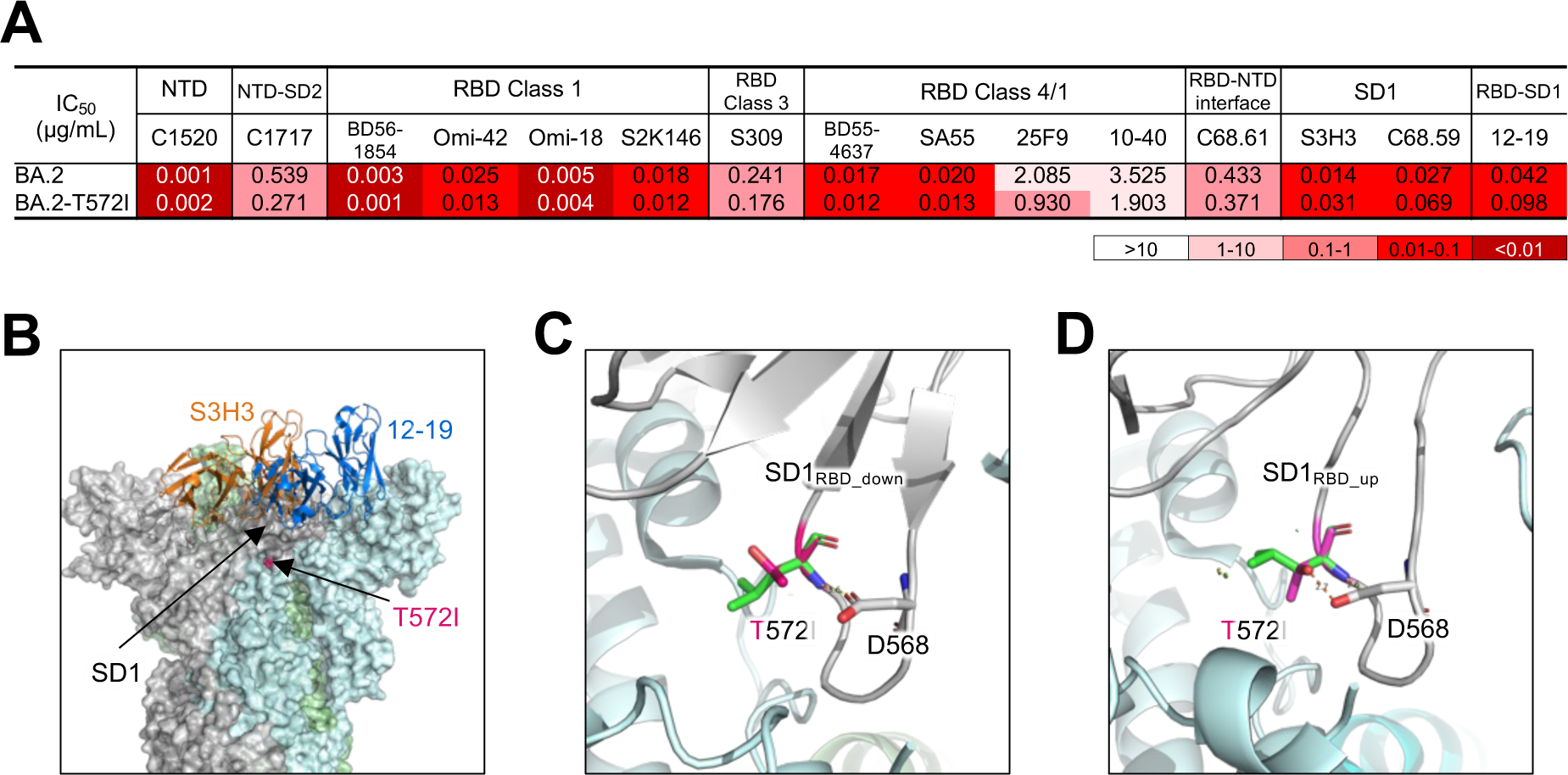
Neutralization of BA.2 and BA.2-T572I by mAbs. **A.** Pseudovirus neutralization IC50 values for mAbs against BA.2 and BA.2-T572I. **B.** T572I is located in the SD1 region of the spike protein and does not directly contact the SD1-directed antibodies S3H3 and 12-19. **C.** Structure modeling of T572I in BA.2 spike (PDB: 7XIX) in all RBD down conformation. **D.** Structure modeling of T572I in RBD up conformation of BA.2 spike (PDB: 7XIW).

**Table S1.**
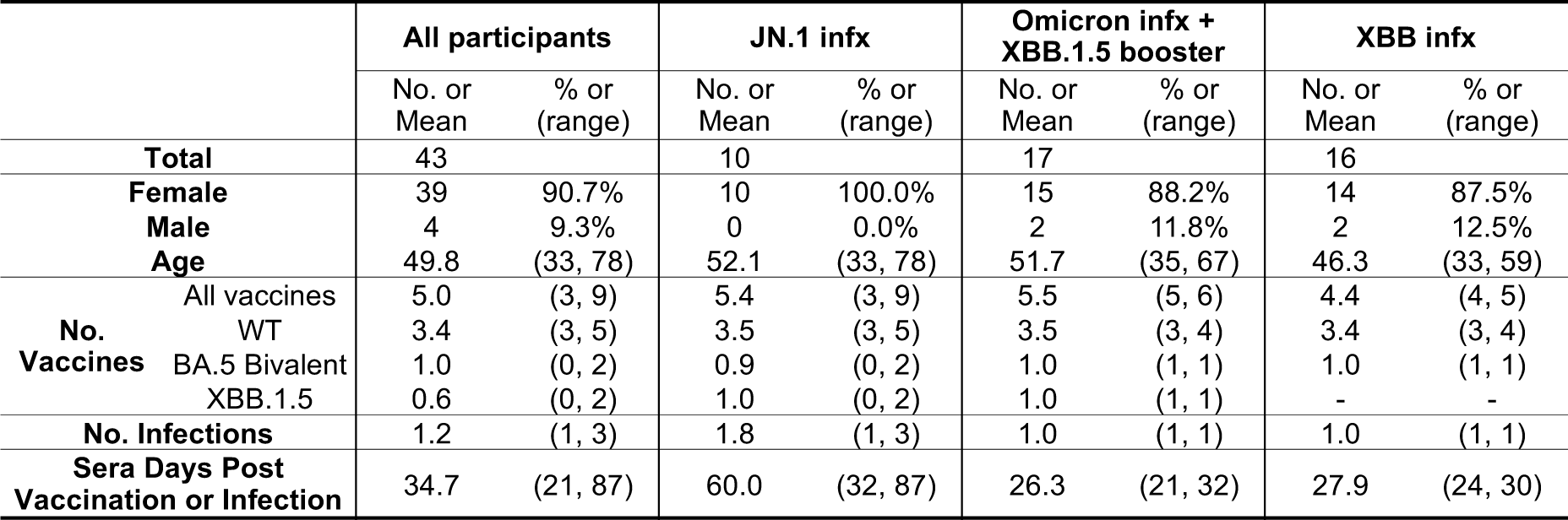
Summary of clinical characteristics of study participants.

**Table S2.** Clinical characteristics of each study participant.

| ID | Group | Age (Yr) | Sex | No. Infx | No. Vax | No. WT Vax | No. BA.5 Bivalent Vax | No. XBB.1.5 Vax | Days Post Infx/Vax | Vaccine History |
| --- | --- | --- | --- | --- | --- | --- | --- | --- | --- | --- |
| 1 | JN.1 infx | 49 | F | 2 | 5 | 3 | 1 | 1 | 35 | WT-P/WT-P/WT-M/BA.5-P/XBB.1.5-P |
| 2 | JN.1 infx | 33 | F | 1 | 4 | 3 | 0 | 1 | 81 | WT-P/WT-P/WT-P/XBB.1.5-P |
| 3 | JN.1 infx | 66 | F | 2 | 6 | 4 | 1 | 1 | 59 | WT-P/WT-P/WT-P/WT-P/BA.5-P/XBB.1.5-M |
| 4 | JN.1 infx | 78 | F | 3 | 6 | 4 | 1 | 1 | 77 | WT-P/WT-P/WT-P/WT-P/BA.5-P/XBB.1.5-P |
| 5 | JN.1 infx | 38 | F | 2 | 5 | 3 | 1 | 1 | 39 | WT-P/WT-P/WT-P/BA.5-P/XBB.1.5-M |
| 6 | JN.1 infx | 35 | F | 2 | 5 | 3 | 1 | 1 | 32 | WT-P/WT-P/WT-P/BA.5-P/XBB.1.5-P |
| 7 | JN.1 infx | 71 | F | 1 | 9 | 5 | 2 | 2 | 80 | WT-M/WT-M/WT-M/WT-M/WT-M/BA.5-M/BA.5-M/XBB.1.5-M/XBB.1.5-P |
| 8 | JN.1 infx | 41 | F | 2 | 5 | 3 | 1 | 1 | 87 | WT-P/WT-P/WT-P/BA.5-M/XBB.1.5-M |
| 9 | JN.1 infx | 48 | F | 1 | 3 | 3 | 0 | 0 | 55 | WT-P/WT-P/WT-P |
| 10 | JN.1 infx | 62 | F | 2 | 6 | 4 | 1 | 1 | 55 | WT-P/WT-P/WT-P/WT-M/BA.5-M/XBB.1.5-P |
| 11 | Om infx + XBB.1.5 vax | 41 | F | 1 | 5 | 3 | 1 | 1 | 25 | WT-M/WT-M/WT-M/BA.5-P/XBB.1.5-M |
| 12 | Om infx + XBB.1.5 vax | 67 | F | 1 | 6 | 4 | 1 | 1 | 29 | WT-P/WT-P/WT-P/WT-P/BA.5-M/XBB.1.5-M |
| 13 | Om infx + XBB.1.5 vax | 52 | F | 1 | 6 | 4 | 1 | 1 | 25 | WT-P/WT-P/WT-P/WT-P/BA.5-P/XBB.1.5-P |
| 14 | Om infx + XBB.1.5 vax | 48 | F | 1 | 5 | 3 | 1 | 1 | 22 | WT-P/WT-P/WT-P/BA.5-M/XBB.1.5-P |
| 15 | Om infx + XBB.1.5 vax | 53 | F | 1 | 5 | 3 | 1 | 1 | 25 | WT-P/WT-P/WT-P/BA.5-P/XBB.1.5-P |
| 16 | Om infx + XBB.1.5 vax | 43 | F | 1 | 5 | 3 | 1 | 1 | 32 | WT-P/WT-P/WT-P/BA.5-M/XBB.1.5-P |
| 17 | Om infx + XBB.1.5 vax | 36 | M | 1 | 5 | 3 | 1 | 1 | 30 | WT-P/WT-P/WT-P/BA.5-M/XBB.1.5-M |
| 18 | Om infx + XBB.1.5 vax | 59 | F | 1 | 6 | 4 | 1 | 1 | 29 | WT-P/WT-P/WT-P/WT-P/BA.5-P/XBB.1.5-M |
| 19 | Om infx + XBB.1.5 vax | 35 | F | 1 | 5 | 3 | 1 | 1 | 29 | WT-P/WT-P/WT-P/BA.5-P/XBB.1.5-P |
| 20 | Om infx + XBB.1.5 vax | 54 | F | 1 | 6 | 4 | 1 | 1 | 22 | WT-P/WT-P/WT-P/WT-M/BA.5-P/XBB.1.5-M |
| 21 | Om infx + XBB.1.5 vax | 58 | F | 1 | 5 | 3 | 1 | 1 | 22 | WT-P/WT-P/WT-P/BA.5-P/XBB.1.5-P |
| 22 | Om infx + XBB.1.5 vax | 61 | M | 1 | 5 | 3 | 1 | 1 | 30 | WT-P/WT-P/WT-M/BA.5-M/XBB.1.5-M |
| 23 | Om infx + XBB.1.5 vax | 42 | F | 1 | 6 | 4 | 1 | 1 | 29 | WT-P/WT-P/WT-P/WT-P/BA.5-M/XBB.1.5-P |
| 24 | Om infx + XBB.1.5 vax | 62 | F | 1 | 6 | 4 | 1 | 1 | 21 | WT-P/WT-P/WT-P/WT-P/BA.5-P/XBB.1.5-P |
| 25 | Om infx + XBB.1.5 vax | 45 | F | 1 | 5 | 3 | 1 | 1 | 22 | WT-P/WT-P/WT-P/BA.5-P/XBB.1.5-P |
| 26 | Om infx + XBB.1.5 vax | 61 | F | 1 | 6 | 4 | 1 | 1 | 26 | WT-P/WT-P/WT-P/WT-P/BA.5-M/XBB.1.5-M |
| 27 | Om infx + XBB.1.5 vax | 62 | F | 1 | 6 | 4 | 1 | 1 | 29 | WT-P/WT-P/WT-P/WT-P/BA.5-P/XBB.1.5-M |
| 28 | XBB infx | 33 | F | 1 | 4 | 3 | 1 | 0 | 30 | WT-P/WT-P/WT-P/BA.5-P |
| 29 | XBB infx | 38 | F | 1 | 4 | 3 | 1 | 0 | 27 | WT-P/WT-P/WT-P/BA.5-P |
| 30 | XBB infx | 59 | F | 1 | 5 | 4 | 1 | 0 | 26 | WT-P/WT-P/WT-P/WT-P/BA.5-P |
| 31 | XBB infx | 44 | M | 1 | 5 | 4 | 1 | 0 | 30 | WT-P/WT-P/WT-P/WT-P/BA.5-M |
| 32 | XBB infx | 54 | F | 1 | 5 | 4 | 1 | 0 | 28 | WT-P/WT-P/WT-P/WT-P/BA.5-P |
| 33 | XBB infx | 38 | F | 1 | 4 | 3 | 1 | 0 | 28 | WT-P/WT-P/WT-M/BA.5-P |
| 34 | XBB infx | 59 | F | 1 | 5 | 4 | 1 | 0 | 30 | WT-P/WT-P/WT-P/WT-P/BA.5-P |
| 35 | XBB infx | 41 | F | 1 | 4 | 3 | 1 | 0 | 29 | WT-P/WT-P/WT-P/BA.5-P |
| 36 | XBB infx | 38 | F | 1 | 4 | 3 | 1 | 0 | 30 | WT-P/WT-P/WT-P/BA.5-P |
| 37 | XBB infx | 42 | M | 1 | 4 | 3 | 1 | 0 | 28 | WT-P/WT-P/WT-P/BA.5-M |
| 38 | XBB infx | 38 | F | 1 | 4 | 3 | 1 | 0 | 24 | WT-P/WT-P/WT-P/BA.5-P |
| 39 | XBB infx | 53 | F | 1 | 4 | 3 | 1 | 0 | 27 | WT-P/WT-P/WT-P/BA.5-P |
| 40 | XBB infx | 40 | F | 1 | 4 | 3 | 1 | 0 | 28 | WT-P/WT-P/WT-P/BA.5-P |
| 41 | XBB infx | 59 | F | 1 | 5 | 4 | 1 | 0 | 24 | WT-M/WT-M/WT-M/WT-M/BA.5-P |
| 42 | XBB infx | 51 | F | 1 | 5 | 4 | 1 | 0 | 29 | WT-M/WT-M/WT-M/WT-M/BA.5-P |
| 43 | XBB infx | 54 | F | 1 | 4 | 3 | 1 | 0 | 29 | WT-P/WT-P/WT-P/BA.5-P |

## References

1. Wang, Q., Guo, Y., Bowen, A., Mellis, I.A., Valdez, R., Gherasim, C., Gordon, A., Liu, L., and Ho, D.D. (2024). XBB.1.5 monovalent mRNA vaccine booster elicits robust neutralizing antibodies against XBB subvariants and JN.1. Cell Host Microbe. 10.1016/j.chom.2024.01.014.

2. Abul, Y., Nugent, C., Vishnepolskiy, I., Wallace, T., Dickerson, E., Holland, L., Esparza, I., Winkis, M., Wali, K.T., Chan, P.A., et al. (2024). Broad immunogenicity to prior SARS-CoV-2 strains and JN.1 variant elicited by XBB.1.5 vaccination in nursing home residents. medRxiv, 2024.2003.2021.24303684. 10.1101/2024.03.21.24303684.

3. Dadonaite, B., Crawford, K.H.D., Radford, C.E., Farrell, A.G., Yu, T.C., Hannon, W.W., Zhou, P., Andrabi, R., Burton, D.R., Liu, L., et al. (2023). A pseudovirus system enables deep mutational scanning of the full SARS-CoV-2 spike. Cell 186, 1263–1278 e1220. 10.1016/j.cell.2023.02.001.

4. Wang, Q., Iketani, S., Li, Z., Liu, L., Guo, Y., Huang, Y., Bowen, A.D., Liu, M., Wang, M., Yu, J., et al. (2023). Alarming antibody evasion properties of rising SARS-CoV-2 BQ and XBB subvariants. Cell 186, 279–286 e278. 10.1016/j.cell.2022.12.018.

5. Kosugi, Y., Kaku, Y., Hinay, A.A., Jr., Guo, Z., Uriu, K., Kihara, M., Saito, F., Uwamino, Y., Kuramochi, J., Shirakawa, K., et al. (2024). Antiviral humoral immunity against SARS-CoV-2 omicron subvariants induced by XBB.1.5 monovalent vaccine in infection-naive and XBB-infected individuals. Lancet Infect Dis. 10.1016/S1473-3099(23)00784-3.

6. Wang, Q., Li, Z., Ho, J., Guo, Y., Yeh, A.Y., Mohri, H., Liu, M., Wang, M., Yu, J., Shah, J.G., et al. (2022). Resistance of SARS-CoV-2 omicron subvariant BA.4.6 to antibody neutralisation. Lancet Infect Dis. 10.1016/S1473-3099(22)00694-6.

7. Wang, Q., Li, Z., Guo, Y., Mellis, I.A., Iketani, S., Liu, M., Yu, J., Valdez, R., Lauring, A.S., Sheng, Z., et al. (2023). Evolving antibody evasion and receptor affinity of the Omicron BA.2.75 sublineage of SARS-CoV-2. iScience 26, 108254. 10.1016/j.isci.2023.108254.

8. Wang, Q., Guo, Y., Zhang, R.M., Ho, J., Mohri, H., Valdez, R., Manthei, D.M., Gordon, A., Liu, L., and Ho, D.D. (2023). Antibody neutralisation of emerging SARS-CoV-2 subvariants: EG.5.1 and XBC.1.6. Lancet Infect Dis 23, e397-e398. 10.1016/S1473-3099(23)00555-8.

9. Qu, P., Xu, K., Faraone, J.N., Goodarzi, N., Zheng, Y.M., Carlin, C., Bednash, J.S., Horowitz, J.C., Mallampalli, R.K., Saif, L.J., et al. (2024). Immune evasion, infectivity, and fusogenicity of SARS-CoV-2 BA.2.86 and FLip variants. Cell 187, 585-595 e586. 10.1016/j.cell.2023.12.026.

10. Ito, J., Suzuki, R., Uriu, K., Itakura, Y., Zahradnik, J., Kimura, K.T., Deguchi, S., Wang, L., Lytras, S., Tamura, T., et al. (2023). Convergent evolution of SARS-CoV-2 Omicron subvariants leading to the emergence of BQ.1.1 variant. Nat Commun 14, 2671. 10.1038/s41467-023-38188-z.

11. Jian, F., Feng, L., Yang, S., Yu, Y., Wang, L., Song, W., Yisimayi, A., Chen, X., Xu, Y., Wang, P., et al. (2023). Convergent evolution of SARS-CoV-2 XBB lineages on receptor-binding domain 455-456 synergistically enhances antibody evasion and ACE2 binding. PLoS Pathog 19, e1011868. 10.1371/journal.ppat.1011868.

12. Juraszek, J., Rutten, L., Blokland, S., Bouchier, P., Voorzaat, R., Ritschel, T., Bakkers, M.J.G., Renault, L.L.R., and Langedijk, J.P.M. (2021). Stabilizing the closed SARS-CoV-2 spike trimer. Nat Commun 12, 244. 10.1038/s41467-020-20321-x.

13. Henderson, R., Edwards, R.J., Mansouri, K., Janowska, K., Stalls, V., Gobeil, S.M.C., Kopp, M., Li, D., Parks, R., Hsu, A.L., et al. (2020). Controlling the SARS-CoV-2 spike glycoprotein conformation. Nat Struct Mol Biol 27, 925–933. 10.1038/s41594-020-0479-4.

14. Li, P., Liu, Y., Faraone, J.N., Hsu, C.C., Chamblee, M., Zheng, Y.M., Carlin, C., Bednash, J.S., Horowitz, J.C., Mallampalli, R.K., et al. (2024). Distinct patterns of SARS-CoV-2 BA.2.87.1 and JN.1 variants in immune evasion, antigenicity, and cell-cell fusion. mBio, e0075124. 10.1128/mbio.00751-24.

15. Kaku, Y., Uriu, K., Kosugi, Y., Okumura, K., Yamasoba, D., Uwamino, Y., Kuramochi, J., Sadamasu, K., Yoshimura, K., Asakura, H., et al. (2024). Virological characteristics of the SARS-CoV-2 KP.2 variant. bioRxiv, 2024.2004.2024.590786. 10.1101/2024.04.24.590786.

16. Li, P., Faraone, J.N., Hsu, C.C., Chamblee, M., Zheng, Y.-M., Carlin, C., Bednash, J.S., Horowitz, J.C., Mallampalli, R.K., Saif, L.J., et al. (2024). Characteristics of JN.1-derived SARS-CoV-2 subvariants SLip, FLiRT, and KP.2 in neutralization escape, infectivity and membrane fusion. bioRxiv, 2024.2005.2020.595020 10.1101/2024.05.20.595020.

17. Khare, S., Gurry, C., Freitas, L., Schultz, M.B., Bach, G., Diallo, A., Akite, N., Ho, J., Lee, R.T., Yeo, W., et al. (2021). GISAID’s Role in Pandemic Response. China CDC Wkly 3, 1049–1051 10.46234/ccdcw2021.255.

18. Smith, D.J., Lapedes, A.S., de Jong, J.C., Bestebroer, T.M., Rimmelzwaan, G.F., Osterhaus, A.D., and Fouchier, R.A. (2004). Mapping the antigenic and genetic evolution of influenza virus. Science 305, 371–376. 10.1126/science.1097211.

19. Wang, Z., Muecksch, F., Cho, A., Gaebler, C., Hoffmann, H.H., Ramos, V., Zong, S., Cipolla, M., Johnson, B., Schmidt, F., et al. (2022). Analysis of memory B cells identifies conserved neutralizing epitopes on the N-terminal domain of variant SARS-Cov-2 spike proteins. Immunity 55, 998–1012 e1018. 10.1016/j.immuni.2022.04.003.

20. Park, Y.J., De Marco, A., Starr, T.N., Liu, Z., Pinto, D., Walls, A.C., Zatta, F., Zepeda, S.K., Bowen, J.E., Sprouse, K.R., et al. (2022). Antibody-mediated broad sarbecovirus neutralization through ACE2 molecular mimicry. Science 375, 449–454. 10.1126/science.abm8143.

21. Nutalai, R., Zhou, D., Tuekprakhon, A., Ginn, H.M., Supasa, P., Liu, C., Huo, J., Mentzer, A.J., Duyvesteyn, H.M.E., Dijokaite-Guraliuc, A., et al. (2022). Potent cross-reactive antibodies following Omicron breakthrough in vaccinees. Cell 185, 2116–2131 e2118. 10.1016/j.cell.2022.05.014.

22. Cao, Y., Jian, F., Wang, J., Yu, Y., Song, W., Yisimayi, A., Wang, J., An, R., Chen, X., Zhang, N., et al. (2022). Imprinted SARS-CoV-2 humoral immunity induces convergent Omicron RBD evolution. Nature 10.1038/s41586-022-05644-7.

23. Pinto, D., Park, Y.J., Beltramello, M., Walls, A.C., Tortorici, M.A., Bianchi, S., Jaconi, S., Culap, K., Zatta, F., De Marco, A., et al. (2020). Cross-neutralization of SARS-CoV-2 by a human monoclonal SARS-CoV antibody. Nature 583, 290–295. 10.1038/s41586-020-2349-y.

24. Cao, Y., Jian, F., Zhang, Z., Yisimayi, A., Hao, X., Bao, L., Yuan, F., Yu, Y., Du, S., Wang, J., et al. (2022). Rational identification of potent and broad sarbecovirus-neutralizing antibody cocktails from SARS convalescents. Cell Rep 41, 111845. 10.1016/j.celrep.2022.111845.

25. Feng, Y., Yuan, M., Powers, J.M., Hu, M., Munt, J.E., Arunachalam, P.S., Leist, S.R., Bellusci, L., Kim, J., Sprouse, K.R., et al. (2023). Broadly neutralizing antibodies against sarbecoviruses generated by immunization of macaques with an AS03-adjuvanted COVID-19 vaccine. Sci Transl Med 15, eadg7404. 10.1126/scitranslmed.adg7404.

26. Liu, L., Iketani, S., Guo, Y., Casner, R.G., Reddem, E.R., Nair, M.S., Yu, J., Chan, J.F., Wang, M., Cerutti, G., et al. (2022). An antibody class with a common CDRH3 motif broadly neutralizes sarbecoviruses. Sci Transl Med, eabn6859. 10.1126/scitranslmed.abn6859.

27. Guenthoer, J., Lilly, M., Starr, T.N., Dadonaite, B., Lovendahl, K.N., Croft, J.T., Stoddard, C.I., Chohan, V., Ding, S., Ruiz, F., et al. (2023). Identification of broad, potent antibodies to functionally constrained regions of SARS-CoV-2 spike following a breakthrough infection. Proc Natl Acad Sci U S A 120, e2220948120. 10.1073/pnas.2220948120.

28. Wang, Q., Guo, Y., Liu, L., Schwanz, L.T., Li, Z., Nair, M.S., Ho, J., Zhang, R.M., Iketani, S., Yu, J., et al. (2023). Antigenicity and receptor affinity of SARS-CoV-2 BA.2.86 spike. Nature. 10.1038/s41586-023-06750-w.

29. Yang, S., Yu, Y., Xu, Y., Jian, F., Song, W., Yisimayi, A., Wang, P., Wang, J., Liu, J., Yu, L., et al. (2024). Fast evolution of SARS-CoV-2 BA.2.86 to JN.1 under heavy immune pressure. Lancet Infect Dis 24, e70-e72. 10.1016/S1473-3099(23)00744-2.

30. Yang, S., Yu, Y., Jian, F., Song, W., Yisimayi, A., Chen, X., Xu, Y., Wang, P., Wang, J., Yu, L., et al. (2023). Antigenicity and infectivity characterisation of SARS-CoV-2 BA.2.86. Lancet Infect Dis 23, e457-e459. 10.1016/S1473-3099(23)00573-X.

31. Zhou, D., Supasa, P., Liu, C., Dijokaite-Guraliuc, A., Duyvesteyn, H.M.E., Selvaraj, M., Mentzer, A.J., Das, R., Dejnirattisai, W., Temperton, N., et al. (2024). The SARS-CoV-2 neutralizing antibody response to SD1 and its evasion by BA.2.86. Nat Commun 15, 2734. 10.1038/s41467-024-46982-6.

32. Hong, Q., Han, W., Li, J., Xu, S., Wang, Y., Xu, C., Li, Z., Wang, Y., Zhang, C., Huang, Z., and Cong, Y. (2022). Molecular basis of receptor binding and antibody neutralization of Omicron. Nature 604, 546–552. 10.1038/s41586-022-04581-9.

33. Liu, L., Casner, R.G., Guo, Y., Wang, Q., Iketani, S., Chan, J.F., Yu, J., Dadonaite, B., Nair, M.S., Mohri, H., et al. (2023). Antibodies targeting a quaternary site on SARS-CoV-2 spike glycoprotein prevent viral receptor engagement by conformational locking. Immunity 56, 2442–2455 e2448. 10.1016/j.immuni.2023.09.003.

34. Wang, Q., Guo, Y., Schwanz, L.T., Mellis, I.A., Sun, Y., Qu, Y., Urtecho, G., Valdez, R., Stoneman, E., Gordon, A., et al. (2024). SARS-CoV-2 Omicron BA.2.87.1 Exhibits Higher Susceptibility to Serum Neutralization Than EG.5.1 and JN.1. Emerg Microbes Infect, 2359004. 10.1080/22221751.2024.2359004.

35. Yisimayi, A., Song, W., Wang, J., Jian, F., Yu, Y., Chen, X., Xu, Y., Yang, S., Niu, X., Xiao, T., et al. (2024). Repeated Omicron exposures override ancestral SARS-CoV-2 immune imprinting. Nature 625, 148–156. 10.1038/s41586-023-06753-7.

36. Kaku, Y., Kosugi, Y., Uriu, K., Ito, J., Hinay, A.A., Jr., Kuramochi, J., Sadamasu, K., Yoshimura, K., Asakura, H., Nagashima, M., et al. (2023). Antiviral efficacy of the SARS-CoV-2 XBB breakthrough infection sera against omicron subvariants including EG.5. Lancet Infect Dis 23, e395-e396. 10.1016/S1473-3099(23)00553-4.

37. WHO (2024). Statement on the antigen composition of COVID-19 vaccines. https://www.who.int/news/item/26-04-2024-statement-on-the-antigen-composition-of-covid-19-vaccines.

38. FDA (2024). POSTPONED - Vaccines and Related Biological Products Advisory Committee May 16, 2024 Meeting Announcement. https://www.fda.gov/advisory-committees/advisory-committee-calendar/postponed-vaccines-and-related-biological-products-advisory-committee-may-16-2024-meeting.

39. Simon, V., Kota, V., Bloomquist, R.F., Hanley, H.B., Forgacs, D., Pahwa, S., Pallikkuth, S., Miller, L.G., Schaenman, J., Yeaman, M.R., et al. (2022). PARIS and SPARTA: Finding the Achilles’ Heel of SARS-CoV-2. mSphere 7, e0017922. 10.1128/msphere.00179-22.

40. Wrapp, D., Wang, N., Corbett, K.S., Goldsmith, J.A., Hsieh, C.L., Abiona, O., Graham, B.S., and McLellan, J.S. (2020). Cryo-EM structure of the 2019-nCoV spike in the prefusion conformation. Science 367, 1260–1263. 10.1126/science.abb2507.

41. Chan, K.K., Dorosky, D., Sharma, P., Abbasi, S.A., Dye, J.M., Kranz, D.M., Herbert, A.S., and Procko, E. (2020). Engineering human ACE2 to optimize binding to the spike protein of SARS coronavirus 2. Science 369, 1261–1265. 10.1126/science.abc0870.

42. Liu, L., Wang, P., Nair, M.S., Yu, J., Rapp, M., Wang, Q., Luo, Y., Chan, J.F., Sahi, V., Figueroa, A., et al. (2020). Potent neutralizing antibodies against multiple epitopes on SARS-CoV-2 spike. Nature 584, 450–456. 10.1038/s41586-020-2571-7.

43. Lefrancios, L., and Lyles, D.S. (1982). The interactionof antiody with the major surface glycoprotein of vesicular stomatitis virus. I. Analysis of neutralizing epitopes with monoclonal antibodies. Virology 121, 157–167.

